# Mechanisms underlying TRPV4-mediated regulation of miR-146a expression

**DOI:** 10.1101/2024.04.03.587984

**Authors:** Bidisha Dutta, Manisha Mahanty, Lakshmyya Kesavalu, Shaik O. Rahaman

**Author notes:** To whom correspondence should be addressed: Shaik O. Rahaman, University of Maryland, Department of Nutrition and Food Science, College Park, MD 20742.

## Abstract

Persistent inflammation is a major contributor in the development of various inflammatory diseases like atherosclerosis. Our study investigates how transient receptor potential vanilloid 4 (TRPV4), a mechanosensitive ion channel, interacts with microRNA-146a (miR-146a), within the context of inflammation and atherosclerosis. Micro-RNAs play a critical role in controlling gene expression, and miR-146a is notable for its anti-inflammatory actions. TRPV4 is activated by diverse soluble and mechanical stimuli, and often associated with inflammatory responses in various diseases. Here, we find that TRPV4 negatively regulates miR-146a expression in macrophages, especially following stimulation by lipopolysaccharides or alterations in matrix stiffness. We show that in atherosclerosis, a condition characterized by matrix stiffening, TRPV4 decreases miR-146a expression in aortic tissue macrophages. We find that TRPV4’s impact on miR-146a is independent of activation of NFκB, Stat1, P38, and AKT, but is rather mediated through a mechanism involving histone deacetylation instead of DNA methylation at the miR-146a promoter site. Furthermore, we show that N-terminal residues 1 to 130 in TRPV4 is essential in suppression of miR-146a expression in LPS-stimulated macrophages. Altogether, this study identifies a regulatory mechanism of miR-146a expression by TRPV4 which may open new potential therapeutic strategies for managing inflammatory diseases.

## INTRODUCTION

Persistent inflammation plays a crucial role in the development of inflammation-driven diseases such as atherosclerosis, foreign body response, and fibrosis (1–7). Macrophages, pivotal in regulating inflammation and tissue repair, possess the capability to detect and eliminate foreign invaders and apoptotic cells due to their phagocytic nature. The adaptability of macrophages supports inflammation management, tissue repair, and regeneration, with their dysregulation potentially leading to numerous pathological states (1–7). The effectiveness of macrophage-driven inflammation hinges on the coordinated expression of key proteins and microRNAs (miRs) involved in their activation, polarization, and phagocytosis, necessitating tight regulation of these factors at transcriptional and post-transcriptional levels through a network of transcription factors (TFs), co-regulators, and epigenetic modifications (1–10). Key TFs such as PU.1, IRF8, STAT1, NFκB, AP1, and PPAR have been extensively studied for their roles in macrophage function (4–7). Additionally, emerging research underscores the significance of mechanical forces in immune response regulation, positioning immune cell mechanotransduction as a growing field (1–3, 11, 12). These research highlights how mechanical cues, present in both normal and diseased states, influence immune cell movement, activation, and inflammatory responses, opening new avenues for understanding and potentially manipulating immune processes.

MiRs, particularly miR-146a, are crucial in regulating macrophage functions and immune response (8–10). Approximately 22 nucleotides in size, these small noncoding RNAs modulate gene expression, and numerous cellular responses (8–10). Notably, miR-146a plays a key role in maintaining homeostasis, and adaptive and innate immunity, primarily by negatively regulating the toll-like receptor (TLR) pathway (8–10, 13, 14). This finding is supported by studies using knockout mice, which showed an increased inflammatory response to endotoxin exposure in the absence of miR-146a (9, 10). Its significant involvement in inflammation also links miR-146a to chronic inflammatory diseases, aging, and malignancies (13, 14). Recent research has expanded to explore miR146a’s role in mechanotransduction, particularly its upregulation in human small airway epithelial cells under oscillatory pressure, and its central role in ventilator-induced lung injury in ARDS patients, through both in vitro and in vivo studies (15, 16).

TRPV4, a member of the TRPV ion channel family, is a nonselective, mechanosensitive channel expressed widely in immune and non-immune cells (17, 18). It responds to various stimuli, including mechanical stretch, osmolarity changes, and biochemical signals like arachidonic acid and cytokines (17, 18). TRPV4 plays significant roles in diverse conditions, such as fibrosis, neuropathies, foreign body responses, and vascular tone regulation (17–23). Research, including our group’s work, highlights TRPV4’s involvement in macrophage activation, polarization, and phagocytosis (17–23). However, its impact on miRs is not well understood. This study utilizes miR sequencing to reveal that miR-146a levels increase when TRPV4 is absent in tissues. Through experiments involving TRPV4-deficient cells and the use of an antagonist, it is shown that TRPV4 inhibits the enhancement of miR-146a expression induced by LPS in macrophages. Furthermore, we show that miR-146a levels decrease in macrophages exposed to high matrix stiffness, as well as in aortic tissues in atherosclerosis-a condition characterized by matrix stiffening- and this decrease is driven by TRPV4. This study further uncovers that TRPV4 influences miR-146a levels through an epigenetic mechanism involving histone deacetylation. Additionally, we demonstrate that the N-terminal residues 1 to 130 of TRPV4 are critical for the suppression of miR-146a expression in macrophages. Overall, these findings highlight a previously unrecognized function of TRPV4 in controlling miR-146a expression.

## MATERIALS AND METHODS

### Reagent and antibodies

The following antibodies were purchased from Cell Signaling Technology, Danvers, MA: p-P65 (cat# 3033S), P65 (cat# 8242S), p-Stat1 (cat# 9167S), Stat1 (cat# 9172S), p-P38 (cat# 9211S), P38 (cat# 9212S), p-JNK (cat# 4668S), JNK (cat# 9252S), p-ERK (cat# 9101S), ERK (cat# 4695S), p-AKT (cat# 9271S), AKT (cat# 4691S), His-H3 (cat# 9715S), Ac-his-H3 (Lys56) (cat# 4243S), and actin (cat# 4970S). Secondary antibodies from Jackson Immunoresearch, including goat, rabbit, and mouse antibodies, were also obtained. For Immunofluorescence-Fluorescent in situ hybridization (IF-FISH) studies antibody for CD68 was purchased from Bio-Rad (cat# MCA1957) and ViewRNA ISH Tissue kit (cat# QVT0600C) was obtained from Thermo Fisher Scientific. ProLong Diamond 4′,6-diamidino-2-phenylindole (DAPI) (cat# P36962) was purchased from Invitrogen. For the calcium influx assay using spinning disc microscopy, we used Calbryte 590 AM from AAT biosciences (cat# 20700). TRPV4 antagonist GSK2193874 (cat# SML0942) was obtained from Sigma Aldrich and agonist GSK 1016790A (cat# 6433) was acquired from Tocris. Pierce RIPA buffer (cat# 89900), and Halt protease and phosphatase inhibitor (cat#78442) was obtained from Thermo Fisher Scientific. Interferon gamma was obtained from R&D Systems, Trichostatin A (cat# T8552) and lipopolysaccharide (LPS) (cat# L5418) was obtained from Sigma Aldrich. Recombinant mouse M-CSF was obtained from R&D Systems. miRNeasy mini kit (cat# 1038703), miRCURY LNA RT kit (cat# 339340), miRCURY LNA SYBR Green kit (cat# 339346) was purchased from Qiagen (Hilden, Germany). iTaq Universal SYBR Green One step RT-PCR kit (cat# 1725150) was purchased from Bio-Rad. Mature miR-146a (cat# YP00204688), pre-miR-146a (cat# YCP2150827) and SNORD68 (cat# YP00203911) primers used for reverse transcription quantitative polymerase chain reaction (RT-qPCR) was obtained from Gene Globe Qiagen (Hilden, Germany). TRAF6 (cat# qMmuCED0046143) and GAPDH (cat# qMmuCED0027497) primers were obtained from Bio-Rad. Petrisoft plates, collagen-coated polyacrylamide hydrogel of 1 kPa (cat# PS100-COL-1-EA) and 50 kPa (cat# PS100-COL-50-EA) stiffness plates, were obtained from Matrigen. We used the following adenovirus constructs generated by Vector Biolabs: Ad (RGD)-mouseTRPV4-WT (Ad-TRPV4-WT), three deletion constructs of different lengths of mouse WT TRPV4 gene (Ad-TRPV4-Δ1-30, Ad-TRPV4-Δ1-130, and Ad-TRPV4-Δ100-130), and Ad-Vec control. Dulbecco’s modified Eagle’s medium (DMEM), heat inactivated fetal bovine serum (FBS), and other cell culture reagents were purchased from Gibco. Antibiotic and antimycotic solution were purchased from Sigma (St. Louis, MO).

### Mice and atherosclerosis model

Dr Makato Suzuki (Jichi Medical University, Togichi, Japan) originally generated TRPV4 KO mice on C57BL/6 background which was then obtained from Dr. David X. Zhang (Medical College of Wisconsin, Milwaukee, WI, USA). ApoE knockout (KO) mice was obtained from the Jackson Laboratory and ApoE:TRPV4 double knockout (DKO) mice on C57BL/6 background was generated by Rahaman lab. Mice were all housed and bred in a temperature and humidity controlled, germ free environment with food and water *ad libitum.* For all the experiments, Institutional Animal Care and Use Committee guidelines were followed and animal protocols were approved by the University of Maryland review committee. For the thioglycolate mouse peritoneal macrophage (thio-MPM) and bone-marrow derived macrophage (BMDM) collection, all mice were chosen to be approximately 6-10 weeks age. For generating the atherosclerotic model, ApoE KO mice and ApoE:TRPV4 DKO mice were subjected to 12 weeks of high-fat diet (HFD) or control chow diet (24, 25). The aortic root areas were removed surgically.

Double Immunofluorescence-FISH assay followed by DAPI staining was employed to study the level of macrophage accumulation and miR-146a expression in these tissues.

### BMDM and thio-MPM isolation, culture, and adenoviral construct transfection

Bone marrow was harvested from mouse femur and incubated with 25 ng/ml MCSF supplemented DMEM for 7-8 days to obtain mature BMDMs (21, 24, 26). Intraperitoneal injection of 2 ml of thioglycolate solution (90000-294; BD Bioscience) was performed and then 96 h post induction thio-MPMs were isolated. Macrophages were cultured in 10% FBS-supplemented DMEM. TRPV4 KO macrophages were seeded on cell culture plates with DMEM, 10% FBS, and transfected with adenovirus TRPV4 constructs or empty vector control (10^6^ plaque-forming units/ml) for 48 h followed by a fresh medium replacement and another 24 h of incubation with complete DMEM. Fresh DMEM containing LPS (100 ng/ml) was added to the cells and maintained for 24 h followed by isolation of miRs to determine the levels of miR-146a.

### MicroRNA (miR) sequencing

To assess the impact of TRPV4 on matrix stiffness-induced miR expression in vivo, we used a subcutaneous biomaterial implantation model in TRPV4 KO mice compared to WT using collagen-coated polyacrylamide (PA) gel discs of rigid (50 kPa) stiffness (19, 20, 22). We selected this stiffness to mimic stiffness of fibrotic and atherosclerotic tissues. Total RNA of each sample was used to prepare the miR sequencing library, which included the following steps: 1) 3’-adapter ligation; 2) 5’-adapter ligation; 3) cDNA synthesis; 4) PCR amplification; and 5) size selection of ∼130-150 bp PCR amplified fragments (corresponding to ∼15-35 nt small RNAs). The libraries were denatured as single-stranded DNA molecules, captured on Illumina flow cells, amplified in situ as clusters and finally sequenced for 51 cycles on Illumina NextSeq per the manufacturer’s instructions. Sequencing was performed on Illumina NextSeq 500 using TruSeq Rapid SBS Kits (#FC-402-4002, Illumina), according to the manufacturer’s instructions. The clean reads that passed the quality filter were processed to remove the adaptor sequence as the trimmed reads. Trimmed reads were alignment to the miRBase pre-miRNAs. miRNA read counts were normalized as tag counts per million miRNA alignments (27, 28).

### Spinning disc confocal microscopy

Fluorescence imaging of macrophages was done using Perkin Elmer spinning disc laser confocal microscope fitted with 20x objective lens. WT and TRPV4 KO macrophages were cultured on 35 mm glass bottom plates. For the calcium influx measurement, the cells were washed with HBBS buffer (Corning, cat# 21023105) twice and incubated with Calbryte 590 AM dye in HBBS for 1 h at 37°C. Following two more washes, transient fluorescence measurement was performed by perfusing the cells with or without LPS (100 ng/ml) in HBBS buffer for 2 mins in the dark followed by perfusion with buffer containing 100 nM GSK 1016790A (GSK101), a known TRPV4 agonist, and fluorescence was measured for 10 mins. To measure the 24 h calcium influx, macrophages were treated with LPS for 24 h. The cells were washed followed by incubation with Calbryte 590 AM for 1 h. These cells were then rewashed to remove the excess Calbryte 590 AM followed by calcium influx measurement for 10 mins in the presence of 100 nM GSK101 in HBBS. The fluorescence intensity was analyzed and quantified using Image J.

### Reverse transcription quantitative polymerase chain reaction (RT-qPCR)

We isolated total RNA from thio-MPMs and BMDMs using miRNeasy kit (Qiagen) following manufacturers protocol. Using miRCURY LNA RT kit (Qiagen), we first generated cDNA, and subsequently determined the expression of mature miR-146a and pre-miR-146a relative to SNORD68 using miRCURY SYBR green kit (Qiagen). The expression of TRAF6 was determined relative to GAPDH using iTaq Universal SYBR Green One step RT-qPCR kit (Bio-Rad) employing the Ct method as described in Bio-Rad RT-qPCR user manual.

### Methylation-specific bisulphite sequencing

The regulatory element of miR-146a gene was carefully evaluated before beginning the assay design. Gene sequences containing the target of interest were acquired from the Ensembl genome browser and annotated and re-evaluated against the UCSC genome browser for repeat sequences including LINE, SINE, and LTR elements. Sequences containing repetitive elements, low sequence complexity, high thymidine content, and high CpG density were excluded from the in-silico design process. For the sample preparation, WT and TRPV4 KO macrophages were treated with LPS (100 ng/ml) and cell pellet samples were lysed based on the total cell count per sample and total volume using M-digestion Buffer (ZymoResearch; Irvine, CA; cat# D5021-9) and 5-10 μl of protease K (20 mg/ml), with a final M-digestion concentration of 2x. The samples were then incubated at 65°C for a minimum of 2 h. Using the EZ-96 DNA Methylation-Direct Kit TM (ZymoResearch; Irvine, CA; cat# D5023) as per the manufacturer’s protocol, 20 μl of the supernatant from the sample extracts were bisulfite modified. The bisulfite modified DNA samples were eluted using M-elution buffer in 46 μl reaction. All bisulfite modified DNA samples were amplified using separate multiplex or simplex PCRs. PCRs included 0.5 units of HotStarTaq (Qiagen; Hilden, Germany; cat# 203205), 0.2 μM primers, and 3 μl of bisulfite-treated DNA in a 20 μl reaction. All PCR products were verified using the Qiagen QIAxcel Advanced System (v1.0.6). Prior to library preparation, PCR products from the same sample were pooled and then purified using the QIAquick PCR Purification Kit columns or plates (cat# 28106 or 28183). Recommended PCR cycling conditions: 95°C 15 min; 45 x (95°C 30s; Ta°C 30 s; 68°C 30 s); 68°C 5 min; 4°C. Test samples were run alongside established reference DNA samples with a range of methylation. They were created by mixing high- and low-methylated DNA to obtain samples with 0, 5, 10, 25, 50, 75, and 100% methylation. The high-methylated DNA is in vitro enzymatically methylated genomic DNA with > 85% methylation. The low-methylated DNA is chemically and enzymatically treated with < 5% methylation. Libraries were prepared and library molecules were purified using Agencourt AMPure XP beads (Beckman Coulter; Brea, CA; cat# A63882). Barcoded samples were then pooled in an equimolar fashion before template preparation and enrichment were performed on the Ion ChefTM system using Ion 520TM & Ion 530TM ExT Chef reagents (Thermo Fisher; Waltham, MA; cat# A30670). Following this, enriched, template-positive library molecules were sequenced on the Ion S5TM sequencer using an Ion 530TM sequencing chip (cat# A27764). FASTQ files from the Ion Torrent S5 server were aligned to a local reference database using the open-source Bismark Bisulfite Read Mapper program (v0.12.2) with the Bowtie2 alignment algorithm (v2.2.3). Methylation levels were calculated in Bismark by dividing the number of methylated reads by the total number of reads. An R-squared value (RSQ) was calculated from the controls set at known methylation levels to test for PCR bias.

### Western blot analysis

For Western blot analysis to detect the levels of MAP kinases, AKT, actin STAT and NFκB levels, WT and TRPV4 KO macrophages were either treated with LPS (100 ng/ml) plus IFN*γ* (20 ng/ml) for 0.5, 1, 2, 6 and 24 h or kept untreated for the control group. For the detection of Histone H3 and acetylated Histone H3 levels, macrophages were pretreated with either 0 nM or 100 nM Trichostatin A (TSA) and then treated with 100 ng/ml of LPS in the presence of TSA for 24 h. The control group remained untreated. Whole cell extracts were separated using 10% SDS-PAGE, and probed with antibodies against p-P65 (1:1000), P65 (1:1000), p-STAT1 (1:1000), STAT1 (1:1000), p-P38 (1:1000), P38 (1:1000), p-JNK1/2 (1:1000), JNK1/2 (1:1000), p-ERK (1:1000), ERK (1:1000), p-AKT (1:1000), AKT (1:1000), Actin (1:2000), Histone H3 (1:1000), and Ac-His H3 (1:1000) respectively. The blots were visualized using HRP-conjugated secondary IgGs and analyzed by UVP BioSpectrum.

### Immunofluorescence-fluorescence in situ hybridization (IF-FISH) assay

IF-FISH studies were performed using ViewRNA Tissue Fluorescence assay kit. 7 μm aortic root tissue sections from ApoE KO and ApoE:TRPV4 DKO mice fed on chow or high-fat diet embedded on glass slides were fixed with pre-chilled 4% paraformaldehyde. Tissue sections were then washed once with PBS and protease digested at 40°C followed by three washes in PBS. Next, we used ViewRNA type 6 target probe set diluted in Probe set diluent QT for the hybridization step for 2.5 h at 40°C in a moisture chamber followed by 1 h amplification with preamplifier and amplifier mix solution provided in the kit. Finally, the probe was labeled using the 25x Label Probe mix (Type 6) diluted in Label Probe diluent QF provided in the kit. Counterstaining of the nuclei was performed by DAPI staining. We next performed immunofluorescence staining on the same sections by incubating with anti-CD68 (1:100) antibody. Anti-Rabbit IgG conjugated with Alexa-Fluor 488 (1:200) were used as secondary antibodies for the immunofluorescence. Fluorescence images were captured, and fluorescence intensity was calculated using ImageJ software and results were presented as integrated density.

### Statistical analysis

All data are presented as mean ± SEM. Statistical comparisons between different groups were conducted using Student’s t-test or one-way ANOVA followed by the Bonferroni test. In the notation, “ns” indicates not significant, * denotes p < 0.05, ** denotes p < 0.01, and *** denotes p < 0.001. All experiments involving animals, both in vivo and in vitro, were carried out in a randomized and blinded manner.

## RESULTS

### MiR-146a expression is upregulated in the absence of TRPV4 in vitro and in vivo

To investigate TRPV4’s impact on miR-146a expression, miR sequencing was performed on skin tissue from WT and TRPV4 KO mice implanted with rigid PA hydrogels. Analysis revealed 85 miRs were upregulated and 212 were downregulated in TRPV4 KO compared to WT (Figs. 1A & B). Notably, miR-146a, known for its anti-inflammatory properties, was approximately 2-fold higher in TRPV4 KO tissues (Fig. 1C). Given TRPV4’s previously identified pro-inflammatory role, these findings suggest TRPV4 may downregulate anti-inflammatory miR-146a, promoting inflammation in response to rigid biomaterial implantation which is linked to increased tissue stiffness (20–24, 29). Macrophages are the predominant cell type in the initiation, maintenance, and resolution of inflammation (1–7). Therefore, we next sought to investigate if the absence of TRPV4 also promoted the expression of miR-146a in macrophages. Calcium influx analysis using spinning disc microscopy demonstrated a functional TRPV4 channel in WT macrophages, evidenced by increased calcium influx upon exposure to LPS, and further increase with TRPV4 agonist, GSK101, but significantly reduced in TRPV4 KO macrophages (Figs. D-G).

**Figure 1.**
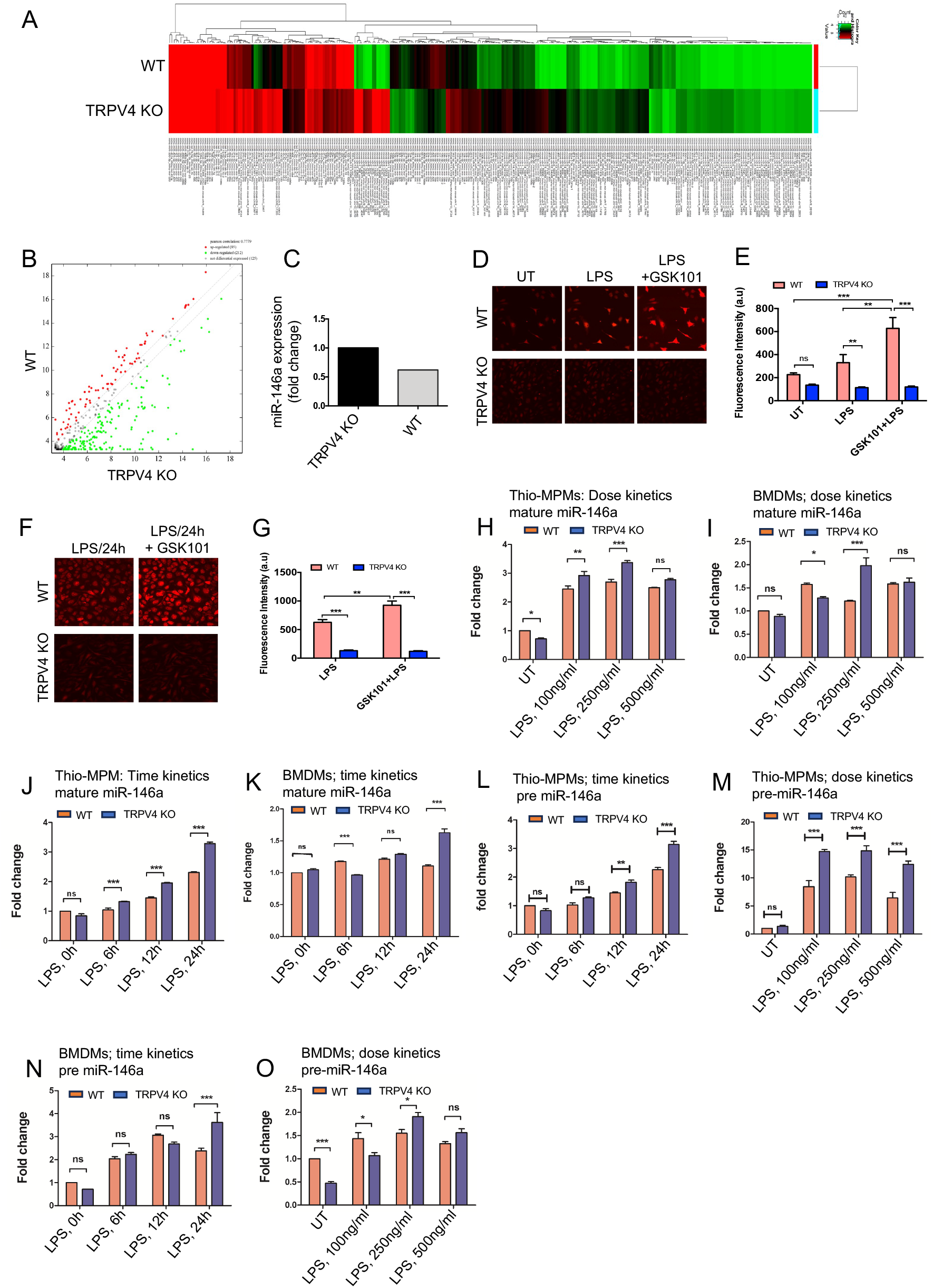
miR-146a shows increased expression levels when TRPV4 is absent, both in vitro and in vivo. **A**. Heatmaps of miRs show differences in expression levels between WT and TRPV4 KO skin tissues implanted with 50 kPa PA hydrogels (n = 5 mice/group; pooled samples). **B**. A scatter plot displays the DEGs based on their log2 fold change values. **C**. Bar graph shows the fold-change in miR-146a expression between WT and TRPV4 KO skin tissues. **D-E**. Spinning disc confocal images reveal calcium influx (red) in WT and TRPV4 KO macrophages under untreated conditions (UT), with LPS (100 ng/ml) alone, and with LPS plus TRPV4-specific agonist, GSK101 (100 nM), treatment (**D**). The quantification of the fluorescence intensity of the cells (n = 50 cells/condition) under all conditions is shown in **E**. **F-G**. Spinning disc confocal images display calcium influx (red) in WT and TRPV4 KO macrophages under LPS treatment, and after LPS treatment for 24 h followed by GSK101 perfusion for 10 mins (**F**). The quantification of the fluorescence intensity of the cells (n = 50 cells/condition) from F is represented in **G**. **H-O**. WT and TRPV4 KO macrophages were treated with 0, 100, 250, and 500 ng/ml of LPS for a dose kinetics study and with 100 ng/ml of LPS for 0, 6, 12, and 24 h for a time kinetics study. The quantification of RT-qPCR analysis shows the expression of miR-146a under different treatment conditions. Plots from RT-qPCR analysis of mature miR-146a expression in macrophages (**H-K**) and pre-miR-146a (**L-O**) under varying doses and time points are presented. Data were analyzed using one-way ANOVA followed by Bonferroni’s multiple comparison test; *p < 0.05, **p < 0.01, ***p < 0.001; ns, not significant.

MiR-146a is known to be a primary response gene in the face of LPS challenge (9). Using LPS to probe miR-146a kinetics in macrophages, we observed a dose-dependent elevation in mature miR-146a expression in WT and TRPV4 KO thio-MPMs, with a notable ∼3-fold increase at 250 ng/ml LPS in WT cells (Fig. 1H). However, TRPV4 KO macrophages exhibited a significantly greater increase in mature miR-146a levels, suggesting TRPV4 modulates miR-146a expression in response to LPS, despite the non-linear response observed in BMDMs (Fig. 1I).

Building on evidence that 100 ng/ml LPS effectively triggers an immune response in macrophages, we selected this concentration for its physiological relevance in subsequent experiments (30). We assessed mature miR-146a expression dynamics in thio-MPMs and BMDMs exposed to 100 ng/ml LPS over 0, 6, 12, and 24 h using RT-qPCR. In WT thio-MPMs, mature miR-146a levels increased steadily over time, doubling after 24 h. TRPV4 KO macrophages exhibited a more pronounced, nearly 4-fold increase in mature miR-146a at the same time point, indicating enhanced expression in the absence of TRPV4 (Fig. 1J). BMDMs displayed a similar pattern, albeit with an irregular increase, confirming the trend in both WT and TRPV4 KO macrophages at 24 h (Fig. 1K).

Investigating whether TRPV4’s influence on miR-146a was post-transcriptional, we analyzed precursor miR-146a (pre-miR-146a) levels in thio-MPMs and BMDMs, observing similar dose and time-dependent responses. WT thio-MPMs displayed a 2.5-fold increase in pre-miR-146a at 24 h and a 10-fold increase at 250 ng/ml LPS (Figs. 1L & M). In TRPV4 knockouts, these increases were even more pronounced, with nearly 3-fold at 24 h and 15-fold at 250 ng/ml LPS, indicating that TRPV4 absence upregulates pre-miR-146a levels without affecting its maturation to miR-146a (Figs. 1L & M). While BMDMs showed varied upregulation, the trend remained consistent, especially notable at 250 ng/ml LPS and 24 h, with TRPV4 deletion amplifying the effect (Figs. 1N & O). This discrepancy between macrophage types may be due to differences in TLR2 and TLR4 levels (31). Altogether, these results suggest a critical role of TRPV4 in regulating both mature and pre-miR-146a levels in macrophages.

### MiR-146a expression is regulated by matrix stiffness in a TRPV4-dependent manner

Exploring TRPV4’s role in miR-146a regulation, we applied the TRPV4 inhibitor, GSK219, to WT thio-MPMs and BMDMs, then measured mature and pre-miR-146a levels by RT-qPCR. GSK219 at 25 μM significantly increased mature miR-146a expression, yielding a 10-fold rise compared to untreated WT and a 5-fold increase over LPS-treated cells, mirroring effects seen with genetic TRPV4 knockdown (Figs. 2A-C). This upregulation was consistent for both miR-146a forms, indicating pharmacological TRPV4 inhibition similarly boosts miR-146a levels. Since miR-146a targets TRAF6, a crucial component of TLR signaling, we assessed TRAF6 expression. RT-qPCR showed TRAF6 was upregulated in LPS-stimulated WT thio-MPMs but markedly reduced in TRPV4 KO macrophages after LPS treatment, underscoring miR-146a’s increase in TRPV4 KO macrophages (Fig. 2D).

**Figure 2.**
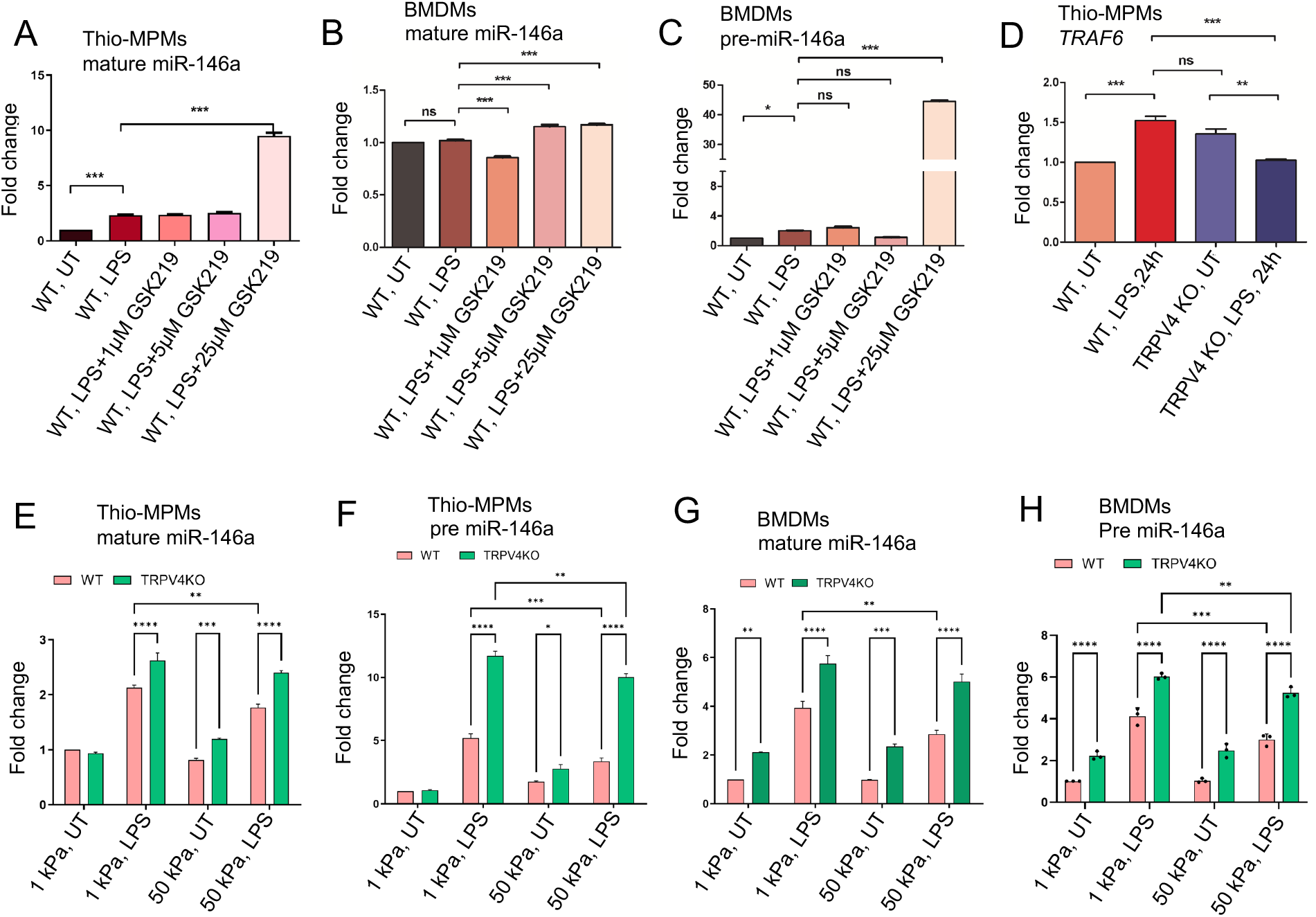
The expression of miR-146a is modulated by matrix stiffness through a process dependent on TRPV4. WT and TRPV4 KO macrophages were either left untreated or treated with LPS (100 ng/ml) alone, or with LPS in combination with 1, 5, or 25 μM of the TRPV4-specific antagonist GSK219. **A-B**. The quantification of RT-qPCR analysis reveals the expression levels of mature miR-146a in thio-MPMs in **A** and in BMDMs in **B**. The expression levels of pre-miR-146a in BMDMs are shown in **C**. **D**. Bar graph displays the fold-change of TRAF6 mRNA in WT and TRPV4 KO macrophages treated with or without 100 ng/ml of LPS for 24 hours. WT and TRPV4 KO macrophages were seeded on 1 and 50 kPa polyacrylamide hydrogel plates coated with collagen (10 μg/ml) and then treated with or without 100 ng/ml of LPS for 24 hours. **E-F**. Bar graphs show the expression levels of mature miR-146a (**E**) and pre-miR-146a (**F**) in thio-MPMs seeded on plates of varying stiffness. **G-H**. The plots display the expression levels of mature miR-146a (**G**) and pre-miR-146a (**H**) in BMDMs seeded on plates of varying stiffness. Three experimental repeats were conducted for each experiment, and data were analyzed using one-way ANOVA followed by Bonferroni’s multiple comparison test; *p < 0.05, **p < 0.01, ***p < 0.001; ns, not significant.

Building on the known mechanosensitivity of miR-146a and TRPV4’s response to matrix stiffness, we explored how TRPV4 influences miR-146a expression in macrophages across different stiffness levels (15, 16, 19, 20). WT and TRPV4 KO thio-MPMs and BMDMs were cultured on PA plates of 1 and 50 kPa, with and without LPS treatment, and miR-146a levels were assessed by RT-qPCR. Both macrophage types showed increased miR-146a expression on softer (1 kPa) compared to stiffer (50 kPa) substrates, particularly under LPS influence (Figs. 2E & F).

The absence of TRPV4 further elevated both pre-and mature-miR-146a levels, indicating TRPV4’s suppressive effect on miR-146a in response to matrix stiffness. This pattern was consistent in BMDMs, suggesting that stiffer environments might suppress miR-146a expression, a process modulated by TRPV4 (Figs. 2G & H).

### N-terminal residues 1 to 130 in TRPV4 is essential in suppression of LPS-induced miR-146a expression in macrophages

The N-terminal domain of TRPV4, known for its role in channel activation by mechanical stimuli like osmolarity and stiffness, was investigated for its potential to regulate miR-146a suppression (20, 32). To this end, three TRPV4 constructs with progressive N-terminal deletions (Ad-TRPV4-Δ1-30, Ad-TRPV4-Δ1-130, and Ad-TRPV4-Δ100-130) were engineered using an adenovirus expression system (Fig. 3A) (20). These constructs, along with the full-length TRPV4 (Ad-TRPV4) and an empty vector (Ad-Vec), were introduced into TRPV4 KO macrophages. Post-LPS treatment, cells expressing the wild-type TRPV4 construct showed reduced miR-146a levels compared to the empty vector control, confirming prior observations (Fig. 3B). This miR-146a suppression was partially relieved in cells transfected with the deletion mutants, indicating the critical role of the TRPV4 1-130 region in mediating LPS-induced miR-146a suppression, a pattern that held even in the absence of treatment, highlighting the importance of the N-terminal domain in miR-146a regulation.

**Figure 3.**
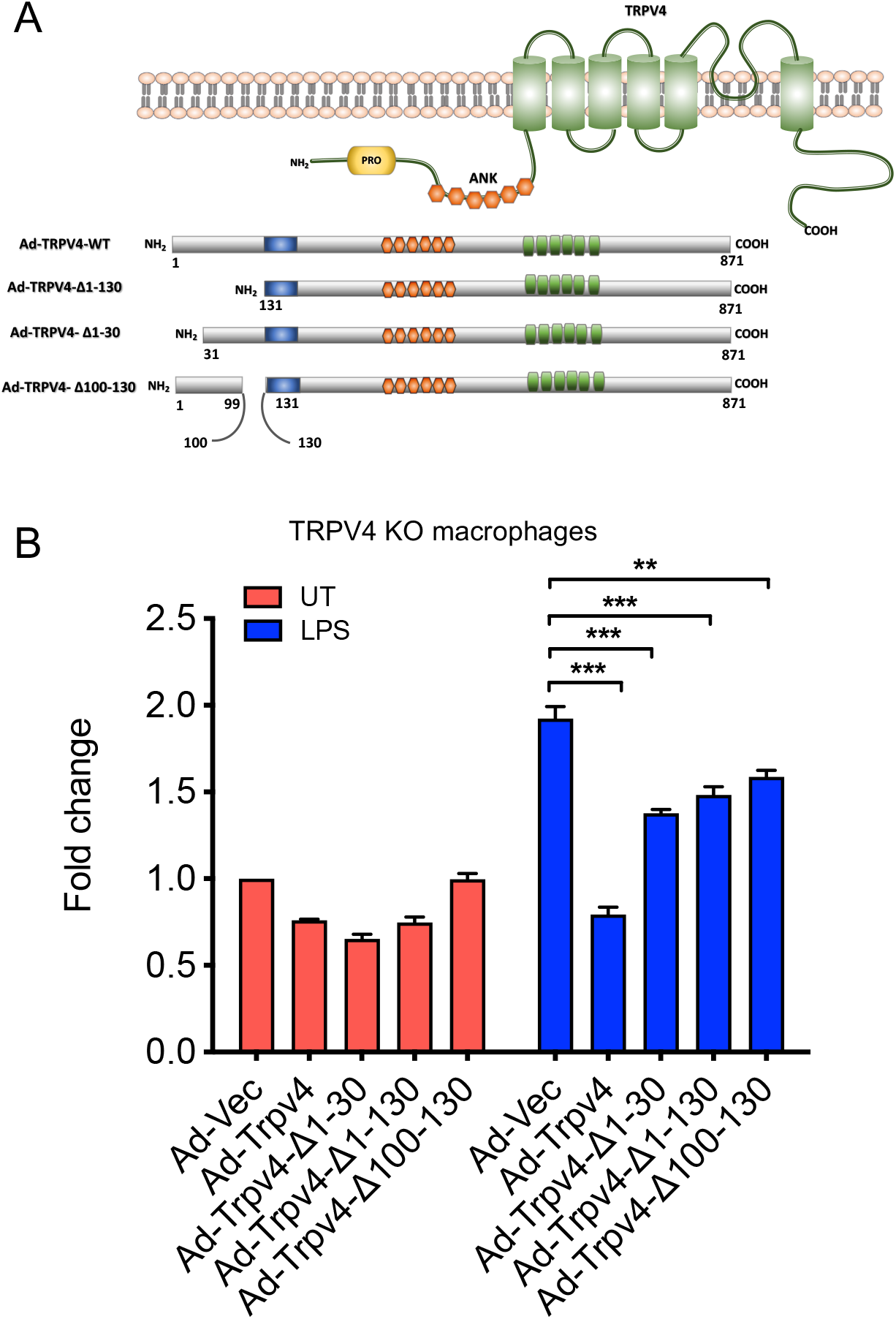
The N-terminal residues 1 to 130 of TRPV4 are crucial for inhibiting LPS-induced miR-146a expression in macrophages. A. The schematic diagram illustrates adenovirus (Ad) expression constructs encoding WT-TRPV4 and three mutant forms with deletions of N-terminal residues 1 to 130, 1 to 30, or 100 to 130. PRD represents the proline-rich domain; ANK denotes ankyrin repeats, alongside the pore domain and the N and C terminals of the TRPV4 protein. **B**. Quantification of RT-qPCR analysis reveals the miR-146a fold-change in TRPV4 KO macrophages transfected with Ad-Vec, Ad-WT-TRPV4, and the three mutant constructs (Ad-TRPV4 del 1-30, Ad-TRPV4 del 1-130, and Ad-TRPV4 del 100-130) under either untreated conditions or after LPS (100 ng/ml) treatment for 24 h. Experiments were repeated three times, and data were analyzed using one-way ANOVA followed by Bonferroni’s multiple comparison test; **p < 0.01, ***p < 0.001.

### TRPV4-regulated suppression of miR-146a is independent of activation of NFκB, Stat1, P38, and AKT

The induction of miR-146a through TLR signaling, known to involve the transcription factor NFκB, which has several binding sites on the miR-146a promoter, was examined in the context of pro-inflammatory stimuli (33). We analyzed the activation of p-P65 in WT and TRPV4 KO macrophages stimulated with LPS and IFNγ, finding no significant difference in its activation levels over time (Figs. 4A & B). Considering recent studies that highlight STAT1’s ability to decrease miR-146a expression by hindering NFκB binding to its promoter (34), our results showed unchanged p-STAT1 levels in both WT and TRPV4 KO macrophages (Figs. 4A & C). Although P38 has been linked to miR-146a expression in endothelial cells (35), our investigations did not reveal its involvement in macrophages, as indicated by stable p-P38 levels (Figs. 4A & D). Further exploration into kinase involvement in miR-146a regulation through TRPV4 highlighted a notable early reduction in p-JNK levels in TRPV4 KO macrophages following stimulation, a change that normalized at later times (Figs. 4A & E). Similarly, p-ERK levels were reduced at 6-and 24-h post-stimulation (Figs. 4A & F). Given the established roles of the JNK and ERK pathways in driving miR-146a expression via transcription factors c-Jun and c-Fos in endothelial cells (35), our findings suggest a nuanced regulatory landscape for miR-146a. Furthermore, our results showed unchanged p-AKT levels in both WT and TRPV4 KO macrophages (Figs. 4A & G).

**Figure 4.**
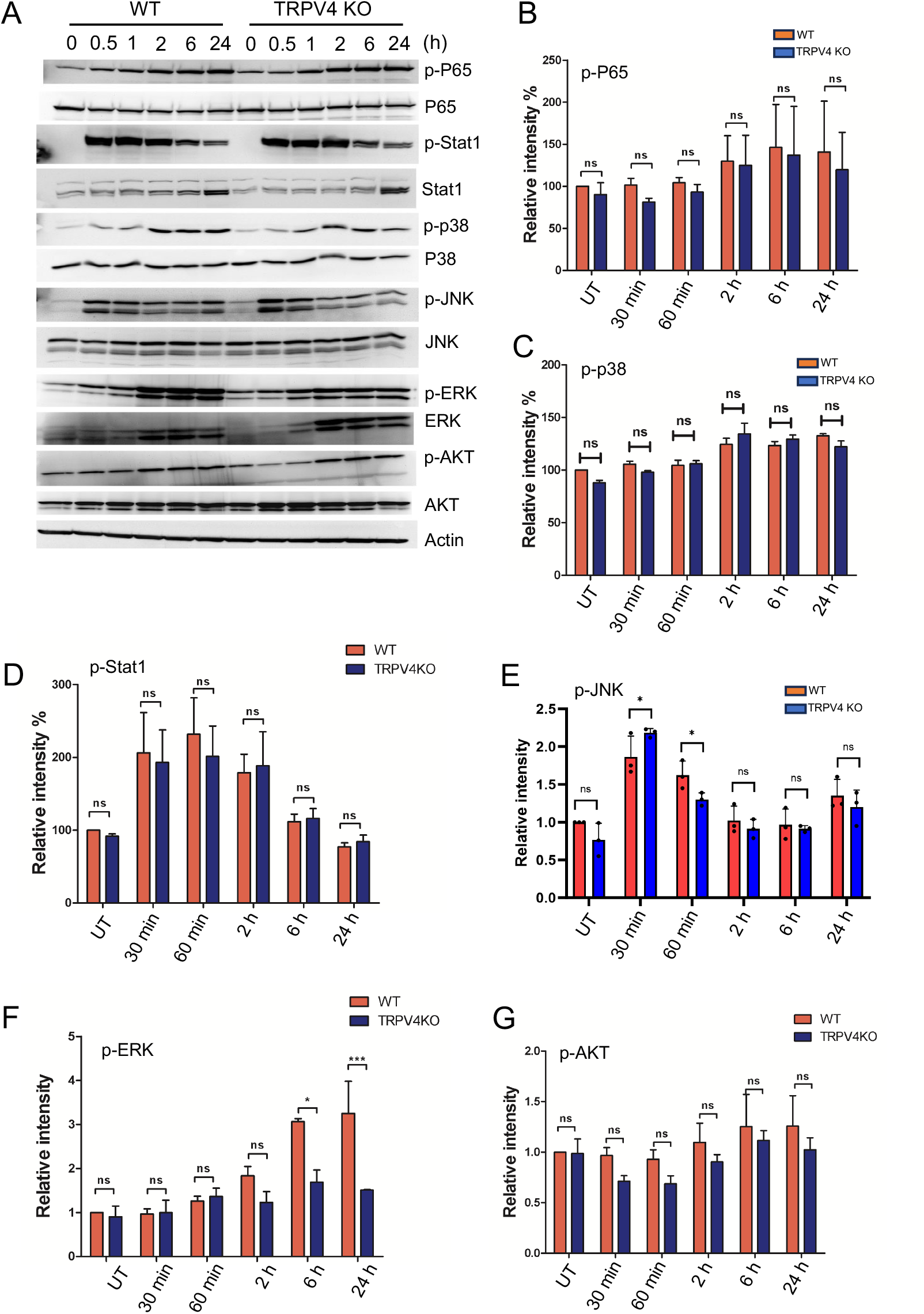
The suppression of miR-146a regulated by TRPV4 occurs independently of the activation of NFκB, Stat1, P38, and AKT. **A**. WT and TRPV4 KO macrophages were treated with LPS (100 ng/ml) plus IFNy (20 ng/ml) for 0.5, 1, 2, 6, and 24 h or kept untreated for the control group. Immunoblotting and quantification of phosphorylated P65, Stat1, P38, JNK1/2, ERK and AKT relative to their total protein levels are shown. Actin was used as loading control. All experiments performed three times and in two biological replicates. **B-G**. Bar graphs show quantification of results from A: p-P65 (**B**), p-P38 (**C**), p-Stat1 (**D**), p-JNK (**E**), p-ERK (**F**), and p-AKT (**G**). Data analyzed using One-way ANOVA followed by Bonferroni’s multiple comparison test; \**p* < 0.05, \*\*\**p* < 0.001; ns, not significant.

### Epigenetic modification is involved in the regulation of miR-146a expression by TRPV4

Over recent decades, epigenetic modifications including the role of miRs in various diseases have been recognized as key regulators of gene expression (36, 37). Given our prior findings showed no significant changes in transcription factor levels, we aimed to explore whether epigenetic alterations could play a role in the TRPV4-mediated suppression of miR-146a. Among such modifications, DNA methylation at CpG sites within gene promoter regions is a prevalent method that affects gene expression. This process does so by altering the accessibility of these regions to transcription factors (38). To determine if TRPV4 influences miR-146a expression through DNA methylation, we first conducted a bioinformatics analysis of miR-146a’s promoter region. This analysis revealed 10 CpG sites located 400 base pairs both upstream and downstream from the transcription start site (Fig. 5A). Given that these sites encompass potential transcription factor bindings and enhancer regions, we proceeded with targeted bisulfite sequencing for these 10 CpG locations. WT and TRPV4 KO macrophages were exposed to 100 ng/ml of LPS for 24 h before undergoing bisulfite sequencing. The sequencing results showed no significant differences in the methylation status of the examined CpG sites between WT and TRPV4 KO cells, indicating that the suppression of miR-146a by TRPV4 may not be mediated by methylation changes at these specific sites (Figs. 5B & C).

**Figure 5.**
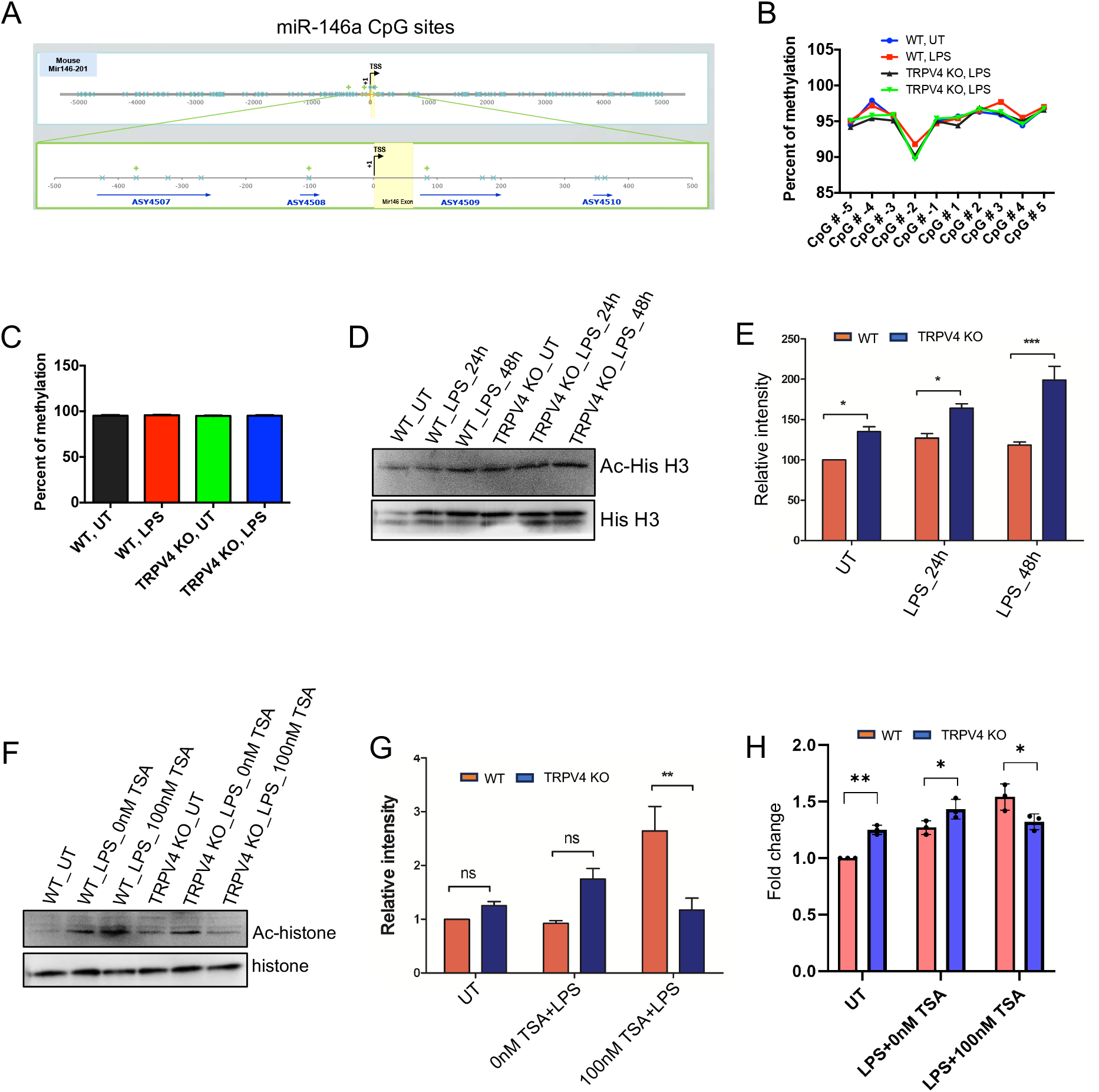
TRPV4 regulates miR-146a expression through epigenetic modification. A. Schematic representation of the transcription start sites of the miR-146a promoter, encompassing the 10 CpG islands (denoted by blue crosses) that were analyzed in our study. Four amplicons were designed to cover the CpG sites of interest, designated by blue arrows. **B**. Percentage of methylation of the 10 CpG sites for conditions WT-UT, WT-100 ng/ml LPS treated (24 h), TRPV4 KO-UT, and TRPV4 KO-100 ng/ml LPS treated (24 h) depicted by a line graph. **C**. Quantification of the percentage of methylation for each condition in the miR-146a gene depicted by a bar plot. **D**. Immunoblotting of acetylated histone H3 relative to total histone levels from WT and TRPV4 KO macrophages treated with or without LPS (100 ng/ml) for 24 and 48 h. **E**. Quantification of the data from D plotted as bar plots, n = 3 independent experiments. **F**. Immunoblot shows acetylated histone H3 in WT and TRPV4 KO macrophages treated with LPS (100 ng/ml) in the presence or absence of the HDAC inhibitor TSA. The control group contained cells that remained untreated (UT). **G**. Quantification of the data from the experiment presented in F shown by a bar plot, n = 3 independent experiments. **H**. Quantification of results from RT-qPCR analysis shows the expression of miR-146a from WT and TRPV4 KO macrophages treated with LPS in the presence or absence of the HDAC inhibitor TSA and in the control untreated (UT) group. Data analyzed by one-way ANOVA followed by Bonferroni’s multiple comparison test; *p < 0.05, **p < 0.01; ns, not significant.

Histone acetylation, a key epigenetic mechanism, promotes gene expression by enhancing chromatin accessibility (39). Research has shown the miR-146a gene’s acetylation, particularly at H3K56, varies under disease states (40). Our study focused on H3K56 acetylation in WT and TRPV4 KO macrophages exposed to LPS over 24 and 48 h, analyzed through Western blot. We observed a notable increase in H3K56 acetylation in TRPV4 KO cells, both under basal and LPS-induced conditions (Figs. 5D & E). Further, treating these cells with the HDAC inhibitor, TSA, with or without LPS, reversed H3K56 acetylation suppression, significantly boosting acetylation in WT cells compared to TRPV4 KO cells (Fig. 5F & G). RT-qPCR was employed to measure miR-146a levels in WT cells compared to TRPV4 KO cells under basal and LPS-stimulated conditions with or without TSA. Results indicated that under basal and LPS-treated conditions, WT macrophages exhibited a suppressed expression of miR-146a compared to TRPV4 KO cells (Fig. 5H). This suppression was alleviated in the presence of TSA showing increased levels of miR-146a in WT compared to TRPV4 cells, indicating that inhibition of deacetylation relieved the suppression on miR-146a expression in WT cells (Fig. 5H). Altogether, this result suggests TRPV4’s role in miR-146a downregulation might be linked to altering H3K56 acetylation, underscoring the epigenetic regulation of gene expression through histone modification.

### TRPV4 suppresses miR-146a expression in atherosclerotic mice model

To validate our in vitro observations in a disease context, we induced atherosclerosis, a chronic inflammatory vascular disease associated with matrix stiffening (41), in ApoE KO mice by administering a high-fat diet (HFD) to create a hyperlipidemic condition. After 12 weeks, aortic root areas from both ApoE KO and ApoE:TRPV4 DKO mice, fed either a chow or HFD, were analyzed through IF-FISH assays. This analysis aimed to assess the effect of TRPV4 knockdown on miR-146a levels within macrophages, utilizing co-labeling with the macrophage marker CD68 to specifically examine macrophage-associated miR-146a expression. Our findings indicate that ApoE KO mice exhibited a significantly higher presence of CD68-positive macrophages under both dietary conditions, suggesting that TRPV4 plays a critical role in macrophage accumulation in both normal and atherosclerotic environments (Figs. 6A & B). Moreover, miR-146a expression was markedly elevated in ApoE:TRPV4 DKO mice across both diets, indicating that TRPV4 deletion leads to an increase in miR-146a levels (Figs. 6A & C). Notably, the proportion of cells positive for both CD68 and miR-146a was greater in the DKO tissues, especially under HFD, highlighting a stronger expression of miR-146a in macrophages lacking TRPV4 (Figs. 6A & D). These findings underscore the significant influence of TRPV4’s absence on macrophage miR-146a levels in inflammatory vascular conditions.

**Figure 6.**
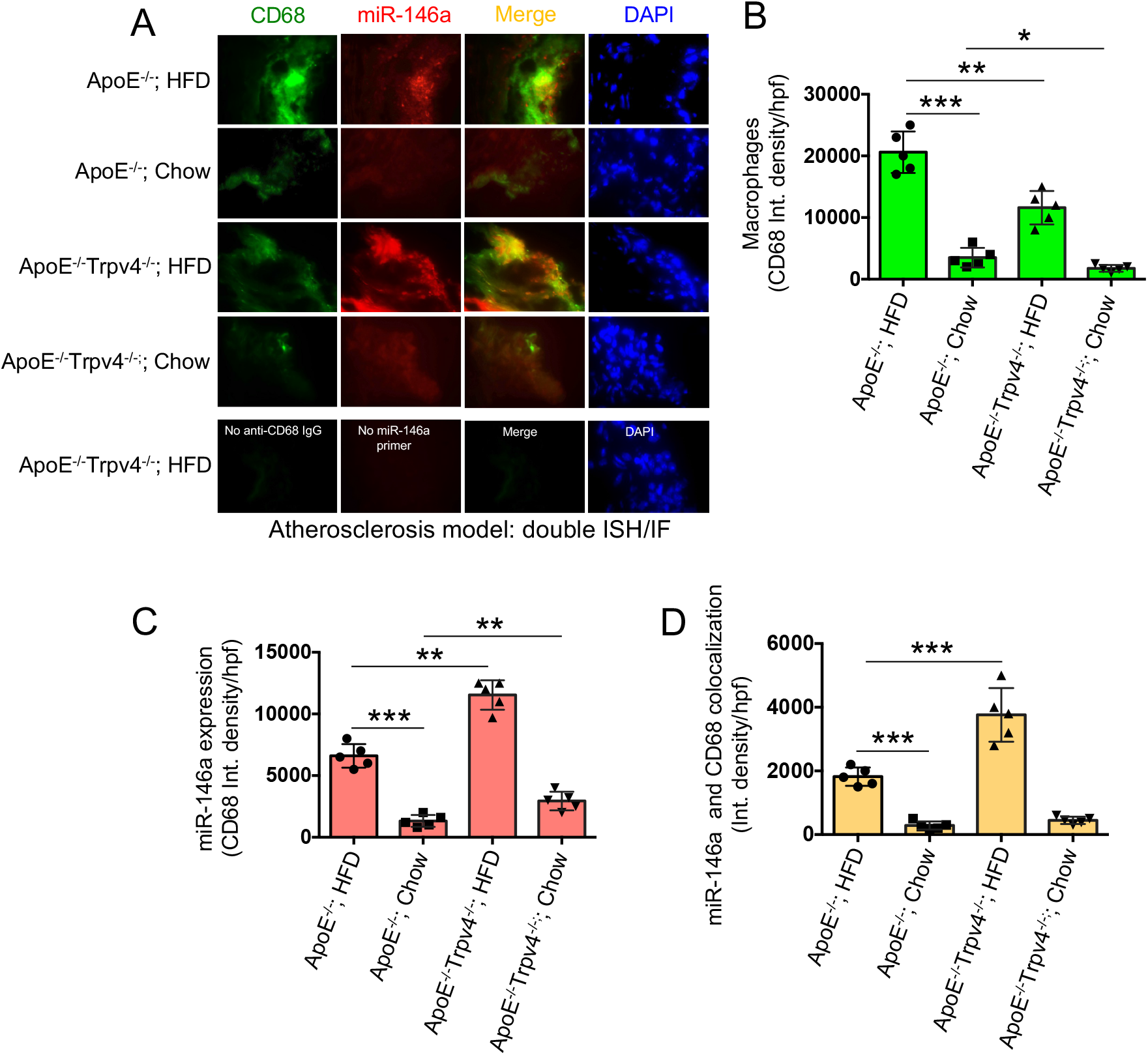
TRPV4 inhibits miR-146a expression in a mouse model of atherosclerosis. Combined fluorescent in situ hybridization (FISH) and immunofluorescence (IF) were carried out on 7 μm aortic root sections from ApoE KO and ApoE:TRPV4 DKO mice fed either chow or a high-fat diet. **A**. Representative IF-FISH images of the tissues hybridized with a miR-146a probe (red), co-stained with a primary antibody for CD68 and an Alexa Fluor conjugated secondary antibody (green). Nuclei were stained with DAPI (blue). Tissues stained in the absence of the miR-146a probe and Alexa Fluor conjugate secondary antibody were used as negative controls. N = 5 mice per group, with 6 images taken for each group. **B**. Quantification of the CD68-positive macrophages in the aortic root tissues under different conditions from figure A. **C**. miR-146a expression quantified for different conditions from figure A. **D**. Quantification of co-stained areas in tissues under different conditions. One-way ANOVA followed by Bonferroni’s multiple comparison test; *p < 0.05, **p < 0.01, ***p < 0.001.

## DISCUSSION

In this study, we found that TRPV4 suppresses miR-146a expression in macrophages, especially after exposure to LPS or due to changes in the matrix stiffness. We show that in atherosclerosis, a chronic inflammatory condition associated with matrix stiffening, TRPV4 decreases miR-146a expression in aortic tissue macrophages. The study demonstrates that the effect of TRPV4 on miR-146a expression in macrophages occurs independently of the commonly implicated signaling pathways such as NFκB, Stat1, P38, and AKT, but rather involves histone deacetylation mechanisms at the miR-146a promoter, not DNA methylation. Additionally, it highlights the importance of TRPV4’s N-terminal residues (1 to 130) in reducing LPS-induced miR-146a levels. Altogether, this study identifies a novel regulatory mechanism of miR-146a expression by TRPV4 which may open new potential strategies for managing inflammatory diseases.

MiR-146a, a key feedback inhibitor of innate immunity, is critical in preventing excessive pro-inflammatory signaling, and its dysregulation is associated with various immune and non-immune inflammatory diseases (8–10, 13–16). Therefore, precise understanding miR-146a’s regulation in both health and disease is vital for developing therapeutic interventions against conditions like atherosclerosis, rheumatoid arthritis, multiple sclerosis, ARDS, and COPD. Soluble factors like LPS trigger miR-146a induction through TLR signaling, a well-documented mechanism (8, 9). Recent research in immune-mechanobiology highlights mechanical factors, such as oscillatory stress, as significant regulators of miR-146a expression, revealing its mechanosensitive role in human lung epithelial cells and implications for lung inflammation (15, 16). Studies in alveolar macrophages and THP1 cells showed miR-146a upregulation in response to such stress, yet the molecular mechanisms behind this regulation remain unclear (15, 16). TRPV4, a mechanosensitive ion channel activated by mechanical stimuli like stiffness and osmolarity, has been implicated in various fibro-inflammatory diseases, where miR-146a dysregulation was also observed (8–10, 19–22, 42–44). Investigating TRPV4’s potential role in miR-146a regulation, our study found upregulation of miR-146a in rigid hydrogel-implanted TRPV4 KO skin tissues compared to WT, suggesting TRPV4’s negative regulatory impact on miR-146a expression. Macrophages, a key cell type in the innate immune system, were utilized to validate sequencing study findings. By utilizing TRPV4 KO macrophages, we demonstrated that TRPV4 deletion leads to a significant increase in miR-146a expression. This connection between TRPV4 activity and miR-146a levels opens new avenues for understanding the molecular mechanisms of miR-146a regulation and its role in inflammatory diseases.

Mechanical ventilation, a treatment for acute respiratory distress syndrome, can induce a pro-inflammatory response and exacerbate lung dysfunction by increasing lung tissue stiffness. Mouse studies revealed that such stiffness leads to elevated miR-146a levels in alveolar macrophages, suggesting mechanical stress influences miR-146a expression (15, 16). However, the exact mechanisms and molecular mediators involved were not fully understood. Our research provides evidence that changes in matrix stiffness directly affect miR-146a levels in primary macrophages, with TRPV4 emerging as a key regulator. Contrary to prior findings linking increased tissue elastance to higher miR-146a expression, we found that higher matrix stiffness suppressed miR-146a compared to lower stiffness levels (45). This discrepancy could be attributed to our use of non-physiological stiffness parameters (50 kPa PA hydrogels), as healthy lung tissue typically exhibits a Young’s modulus between 1.4 and 3.4 kPa (45). Further, our adenoviral overexpression studies confirmed TRPV4’s role in miR-146a regulation, specifically highlighting the importance of its N-terminal residues (1–130) in this process, consistent with their known involvement in TRPV4-mediated calcium influx in macrophages (20).

Studies indicates post-transcriptional regulation of miR-146a among various miRs in T cells (46). Our findings reveal that the expression levels of pre-miR-146a and mature miR-146a in macrophages from both WT and TRPV4 KO mice are comparable, suggesting TRPV4 does not affect the processing from precursor to mature miR-146a. Despite miR-146a being a known NFκB-responsive gene, particularly under LPS stimulation (9), our data show that TRPV4’s influence on miR-146a expression does not involve changes in NFκB’s activity, as indicated by the unchanged phosphorylation status of the P65 subunit. Furthermore, despite recent discoveries of STAT1’s role in miR-146a regulation, our study does not support STAT1’s involvement in TRPV4’s regulatory mechanisms (47). In human umbilical vein endothelial cells, it has been demonstrated that miR-146a expression is regulated through the activation of p38, JNK, and ERK pathways (35). However, our research found no notable differences in the activity of p38, JNK, or AKT pathways between WT and TRPV4 KO cells. However, we observed an increase in phosphorylated ERK in LPS-stimulated WT macrophages, hinting at a potential ERK-mediated pathway for TRPV4 to regulate miR-146a expression, a theory that requires further exploration to substantiate.

Recent research underscores the role of epigenetic modifications in regulating miR-146a, with findings of histone deacetylation suppressing miR-146a in macrophages from aged mice, thereby exacerbating inflammatory responses (14). It’s established that active miR-146a promoters correlate with acetylated histone H3 levels (48). Our investigation into TRPV4 knockout macrophages showed an increase in global histone H3 acetylation, hinting at TRPV4’s potential in modulating H3 acetylation downwards and affecting miR-146a expression. Thus, our study introduces an epigenetic function for TRPV4, expanding upon previous knowledge of TRPA1’s role in epigenetic regulation (49), and positioning TRPV4 as a novel player in the epigenetic modulation of miR-146a.

The remodeling of the extracellular and intracellular matrix can alter arterial tissue stiffness, potentially initiating molecular pathways that contribute to atherogenesis (3, 21, 41). There’s growing evidence linking miR-regulating genes to the onset and progression of atherosclerosis, with miR-146a’s role being particularly debated (50). While some studies have identified an increase in miR-146a within human atherosclerotic plaques and a protective effect against atherosclerosis in hyperlipidemic mice via ApoE-mediated upregulation, others report a pro-atherogenic effect where miR-146a absence reduced atherosclerosis (51–53). We found elevated miR-146a levels in hyperlipidemic ApoE KO mice on a high-fat diet, aligning with observations of miR-146a induction by dietary fats in adipose tissues (54). Further, TRPV4 deletion in ApoE KO mice amplified miR-146a levels in aortic tissues, supporting its pro-atherogenic role through miR-146a modulation. These findings hint at a complex interaction between atherosclerosis, TRPV4, and miR-146a, underscoring the potential of TRPV4 as a target in atherosclerosis management by modulating miR-146a levels.

Our study has notable limitations, including the use of a global TRPV4 KO model, which might activate compensatory mechanisms affecting miR-146a expression due to TRPV4’s wide presence across various cell types. We didn’t explore beyond histone H3’s involvement or investigate the ERK pathway’s potential influence on miR-146a levels, nor did we examine possible interactions between epigenetic modifications and signaling pathways. Despite these constraints, our findings highlight miR-146a as a miR responsive to mechanical stiffness, regulated by the mechanosensitive TRPV4 channel through epigenetic mechanisms under conditions of inflammatory stress.

## ACKNOWLEDGEMENTS

This work was supported by an NIH (R01EB024556) grant to Shaik O. Rahaman. We acknowledge the use of ChatGPT 4 for proofreading the final draft in March, 2024.

## AUTHOR CONTRIBUTIONS

SOR and BD conceived the study, designed, and performed the experiments, and analyzed data. SOR and BD wrote the manuscript. MM assisted with experiments. LK analyzed data and edited the MS. All authors reviewed the MS and approved the final content of the MS.

## DATA AVAILABILITY

All data generated or analyzed during this study are included in this article.

